# Pervasive, conserved secondary structure in highly charged protein regions

**DOI:** 10.1101/2023.02.15.528637

**Authors:** Catherine G. Triandafillou, Rosalind Wenshan Pan, Aaron R. Dinner, D. Allan Drummond

## Abstract

Understanding how protein sequences confer function remains a defining challenge in molecular biology. Two approaches have yielded enormous insight yet are often pursued separately: structure-based, where sequence-encoded structures mediate function, and disorder-based, where sequences dictate physicochemical and dynamical properties which determine function in the absence of stable structure. Here we study highly charged protein regions (>40% charged residues), which are routinely presumed to be disordered. Using recent advances in structure prediction and experimental structures, we show that roughly 40% of these regions form well-structured helices. Features often used to predict disorder—high charge density, low hydrophobicity, low sequence complexity, and evolutionarily varying length—are also compatible with solvated, variable-length helices. We show that a simple composition classifier predicts the existence of structure far better than well-established heuristics based on charge and hydropathy. We show that helical structure is more prevalent than previously appreciated in highly charged regions of diverse proteomes and characterize the conservation of highly charged regions. Our results underscore the importance of integrating, rather than choosing between, structure- and disorder-based approaches.

## Introduction

In the overarching quest to understand how genotype shapes phenotype, the question of how protein sequence encodes protein function has proved a rich and enduring challenge. An early and still pervasive conceptual framework in which stable sequence-encoded protein structures confer biological function has been met by a newer (yet by now firmly established) companion approach born from the recognition that many functions—binding, selective recruitment, formation of large-scale structures, and more—can be achieved by sequences which do not adopt a stable conformation (intrinsically disordered regions, or IDRs). Although neither approach is exclusive of the other, and indeed they anchor a continuum [1], for historical and methodological reasons many analyses adopt one approach or the other based on various heuristics [2–7]. Such heuristics have had an outsized impact on how sequence-function maps are explored.

In one early and influential study of IDRs, Uversky and colleagues discovered that plotting mean net charge against mean hydropathy (hydrophobicity) permits a dividing line to be drawn separating folded from disordered proteins [2,8]. In these analyses, highly charged, weakly hydrophobic sequences have a strong tendency to be disordered. More recently developed heuristics go beyond composition: simulation studies suggest that the degree of mixing of opposite charges within a highly charged, nearly net-neutral (polyampholyte) sequence is a predictor for the biophysical properties of such polypeptides, specifically whether they form expanded or compact structures in solution [3,9]. These studies assume that the sequences in question do not take on well-defined structures, largely based on the observation that many disordered proteins are polyampholytes (∼75% of known IDRs have a fraction of charged residues (FCR) > 0.35 [10]). These findings, although based on a few hand-picked sequences from different organisms and proteins, have broadly informed the analysis of many other sequences [11–13].

Other analyses, while still converging on the general finding that disorder is associated most strongly with a bias toward charged residues and away from hydrophobic residues, have emphasized extreme compositional biases themselves as strong predictors of disorder [4,5]. The most common quantitative description of sequence compositional bias is the Shannon entropy, often referred to as complexity [6,14]. Complexity here has a statistical, not biological, interpretation; “simple” sequences such as homopolymers or sequences composed of only a few types of amino acids have low sequence entropy and thus low complexity.

Low-complexity regions (LCRs) have in the past decade experienced a surge of attention, driven by the observation that they are associated with mesoscale organization in cells: clusters, granules, hydrogels [15–17], membraneless organelles, and a host of related structures now referred to as biomolecular condensates [18]. Charged LCRs in particular play a crucial role in biomolecular condensation in highly influential model systems, mediating complex coacervation [7] and phase separation [19–22]. Particular sequence features such as enrichment with positively charged residues like arginine and with conformationally flexible glycine, most memorably in the RGG motif [23], appear often in RNA binding proteins that are known to condense; interactions with cationic residues can be modulated by negatively charged regions, leading to the proposal of a molecular grammar for such interactions [24].

Together, these lines of inquiry both reflect and create conditions in which highly charged, low-hydrophobicity LCRs may be studied nearly exclusively through the lens of disorder [3,10]. Because of the historical roots of the disorder presumption—particularly that many of the paradigm-shaping observations were made as databases of sequences and structures were in their infancy—the presumption itself has persisted with few challenges.

A confluence of trends and events has laid the groundwork for a productive reexamination of these assumptions. First, the maturation of structural and sequence databases has prompted increasingly critical looks at our understanding of LCRs [25] and IDRs [26]. In parallel, specific examples have accumulated of well-defined structure in sequences which would, by existing heuristics, be overwhelmingly predicted to be disordered: alpha-helices in myosin [27] and caldesmon [28], and a coiled-coil region in the mRNA export protein GLE1 [29]. Indeed, there is a long-established connection between charge patterning and helix formation [10–13,30,31]. Finally, new methods now permit more reliable and farther-reaching assessment of the disorder presumption for highly charged regions, notably high-quality structure prediction [32,33].

So motivated, we return to the root issues: to what extent are naturally occurring highly charged protein regions structured versus disordered? What is the empirical relationship between the fraction and patterning of charged residues and the biophysical properties of a region? Are highly charged regions conserved over evolutionary time, as we would expect for biologically important properties? And how easily can one distinguish between structured and disordered regions on the basis of simple sequence heuristics?

To answer these questions, we systematically identify highly charged regions proteome-wide in related eukaryotes (budding yeasts) and characterize their sequence properties, predicted structure, and evolution. In contrast to previous studies [3,9,10,34], our work examines the entire proteome, permitting us to quantify the frequency and, with proteome-scale homology, evolutionary conservation of charged regions at the genomic scale. We find that naturally occurring polyampholytes are highly prevalent and, despite being low-complexity with often-poor sequence conservation, these regions are often predicted to form or contain alpha-helices. We confirm these predictions through comparison with experimentally derived structures. These results demonstrate that certain LCRs, even those enriched for charged amino acids thought to be important for intermolecular interactions driving phase separation, may adopt a well-defined structure in the right sequence or physicochemical context. More broadly, we show that it is important to consider structural properties explicitly when evaluating the other properties of an LCR.

## Results

### Highly charged regions of low sequence complexity are prevalent in the yeast proteome

We first established criteria for regions to be “highly charged” and used them to identify regions in the *S. cerevisiae* proteome, our departure point owing to its extensive experimental and evolutionary characterization. Examining amino acid usage (Figure 1a), we found that the charged residues (glutamate, aspartate, lysine, and arginine) together constitute 23% of the total amino acids. Unlike all other categories of amino acids, the frequency of each charged amino acid (as determined from the frequency of observed codons) deviates strongly from expectation based on the underlying nucleotide frequency (Figure 1a, light gray points), evidence for evolutionary selection.

**Fig. 1.**
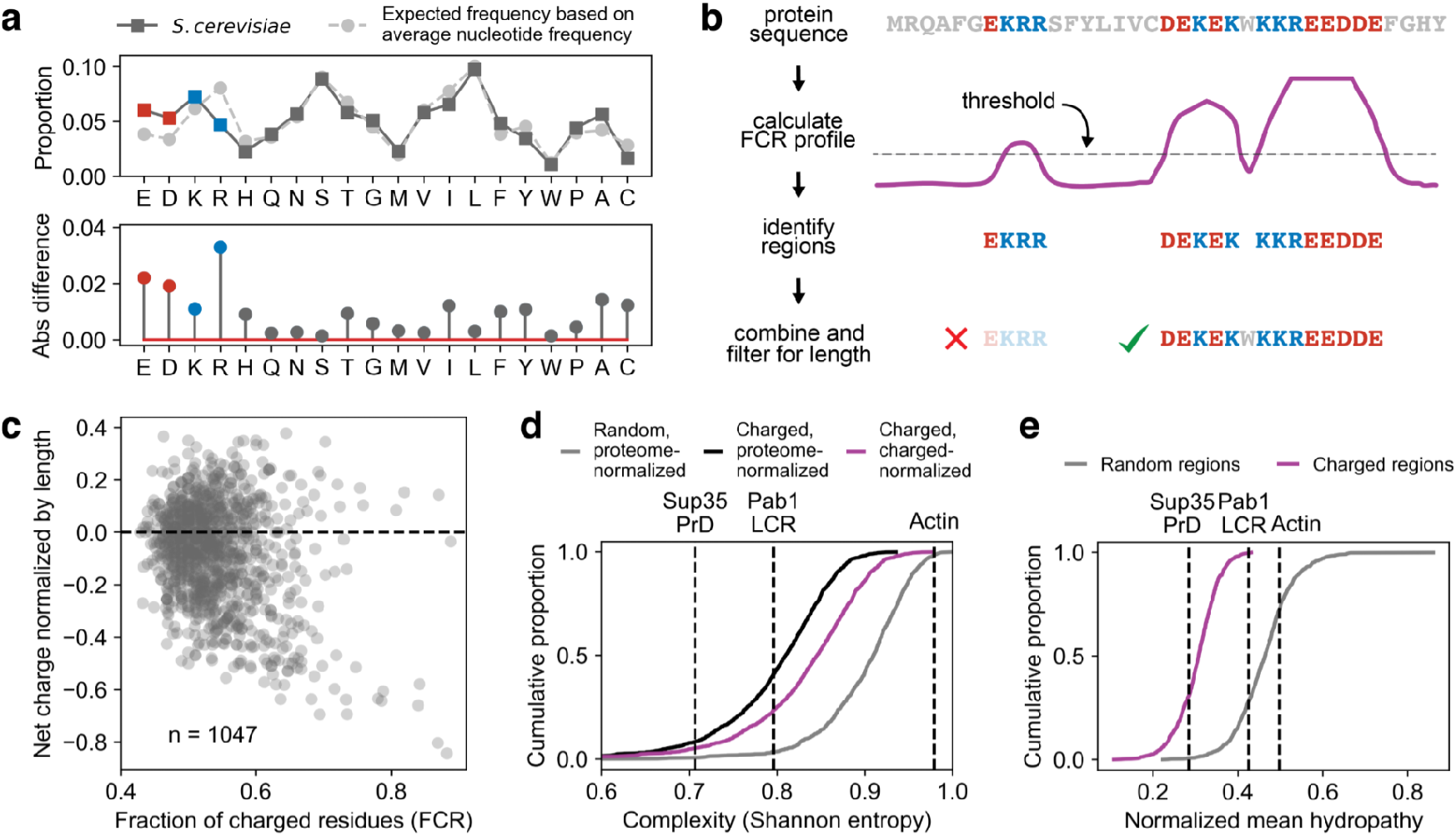
Regions high in charge are prevalent in the yeast proteome and are low complexity. **a** Top: the average frequency of each amino acid in the yeast proteome. Light gray trace is the expected frequency based on the average nucleotide frequency and the codons which code for each amino acid. Bottom: The difference between the dark and light traces in the top panel. **b** Cartoon of the algorithm which detects highly charged regions. **c** Summary of the per-region fraction of charged residues (FCR) and normalized net charge for the regions detected by the algorithm shown in **b. d** Complexity of the charged regions. The complexity was calculated according to the compositional entropy defined in [14]. Reference sequences are two known low complexity domains from Sup35 (a typical prion-like low-complexity protein) and Pab1 (atypical hydrophobic-rich intrinsically disordered region), and a high complexity folded sequence (actin). **e** Normalized hydropathy of the charged regions (purple) and length-matched randomly drawn regions (gray). The reference sequences are the same as those in **d**.

To isolate highly charged regions, we took a sliding-window approach, moving a 12-amino-acid window across all protein sequences and selecting regions with a fraction of charged residues (FCR) above 0.4, with some tolerance for transient deviations (see Figure 1b and Methods). After trimming uncharged ends off these segments, the resulting highly charged regions have a median length of 50 and a FCR ≥ 0.43 (Figure 1c and S1c), more stringent than, for example, a published definition of a strong polyampholyte (FCR > 0.30) [35]. We identified 1,047 regions in 800 proteins; about 14% of protein-coding genes encode at least one highly charged region. The FCR in these regions is just over two standard deviations above that for randomly chosen regions in the proteome (Figure S1a); the regions also have substantially higher charge density than the proteins which contain them (Figure S1b).

We examined the distribution of both FCR and normalized net charge across all regions; it is more common for a region to have a net charge close to zero, although there is a significant number of net negatively charged regions (Figure 1c). Examples of neutral regions and those that carry a net charge can be found in Table 1.

**Table 1:**
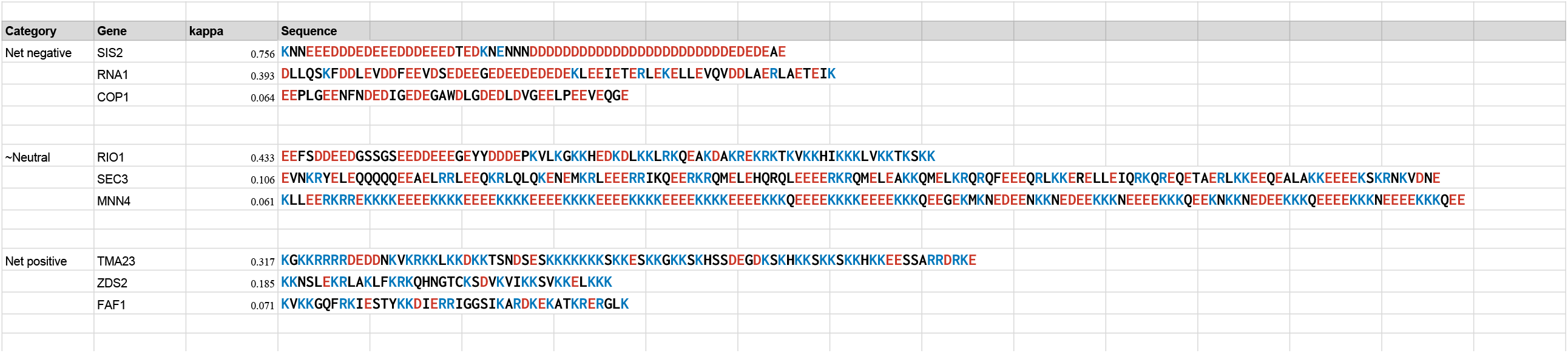
Example highly charged regions from the yeast proteome.

We expected that the highly charged regions would have lower complexity and hydrophobicity than random regions, because they must be enriched for a small subset of (charged) amino acids, and we confirmed that this is the case (Figure 1d,e). To determine the regions’ complexity relative to our selection bias, we calculated the complexity normalized by the entropy of a mostly charged proteome (50% charged amino acids and all other amino acids equally represented, see Methods for details, Figure 1d purple trace). Even with this correction, the regions are far less complex than randomly drawn regions (P < 10^−6^, Wilcoxon rank sum test), suggesting a further bias in amino acid usage in these regions beyond that explained by their enrichment for charge.

We also examined the distribution of the proteins containing these regions within the cell. We found that they were enriched in the nucleus, and especially in the nucleolus (Figure S1d), consistent with recent findings that across several species nucleolar proteins are enriched for charge-rich low-complexity sequences [36].

In summary, we find that highly charged regions which exceed even stringent definitions of polyampholytes are common in the yeast proteome, are on average less hydrophobic and less complex than average sequences, and are enriched in specific nuclear compartments.

### Secondary structure is pervasive in highly charged regions

Given the historical and intuitive associations between low hydrophobicity/high charge density and disorder, we predicted that the vast majority of the regions we identified would not adopt a well-defined structure. We thus set out to assess what proportion of the regions we identified were IDRs using experimentally derived structures and recently available proteome-wide structure prediction (AlphaFold) [32]. Although the biophysical properties of disordered regions cannot be accurately assessed using AlphaFold structures [37], disorder can be inferred in two ways. The first is to impute disorder to residues with a low AlphaFold confidence score [37]; the second is to ascribe disorder to regions with high-confidence coil (e.g., not helix or sheet) predictions. We employ both methods. To assess the validity of these choices, we analyzed the predicted AlphaFold structure of protein regions from the DisProt database (which contains proteins that have been empirically measured to be disordered through a variety of experimental means including circular dichroism and NMR) and found that the vast majority of confidently predicted residues in these regions were scored as “coil” by DSSP (Figure S2a). Thus, by using AlphaFold we were able to assess both structure and disorder—with the same method—proteome-wide.

Returning to the highly charged regions, we used the AlphaFold predictions to classify each residue as either disordered (low confidence, or high-confidence and scored as coil) or ordered (high-confidence and scored as helix or sheet; see Methods for a complete description of scoring cutoffs). While a significant number of the highly charged regions were almost completely composed of residues classified as disordered, in many regions (40% of the total) more than half of the residues were predicted to be structured (Figure 2a). We examined the secondary structure classification for all confidently predicted residues (45% of the total) across the entire dataset. The highly charged regions were markedly enriched for alpha-helical secondary structure compared to disordered regions from Disprot and had a similar frequency to length-matched randomly drawn regions (Figure 2b, Figure S2a).

**Fig. 2.**
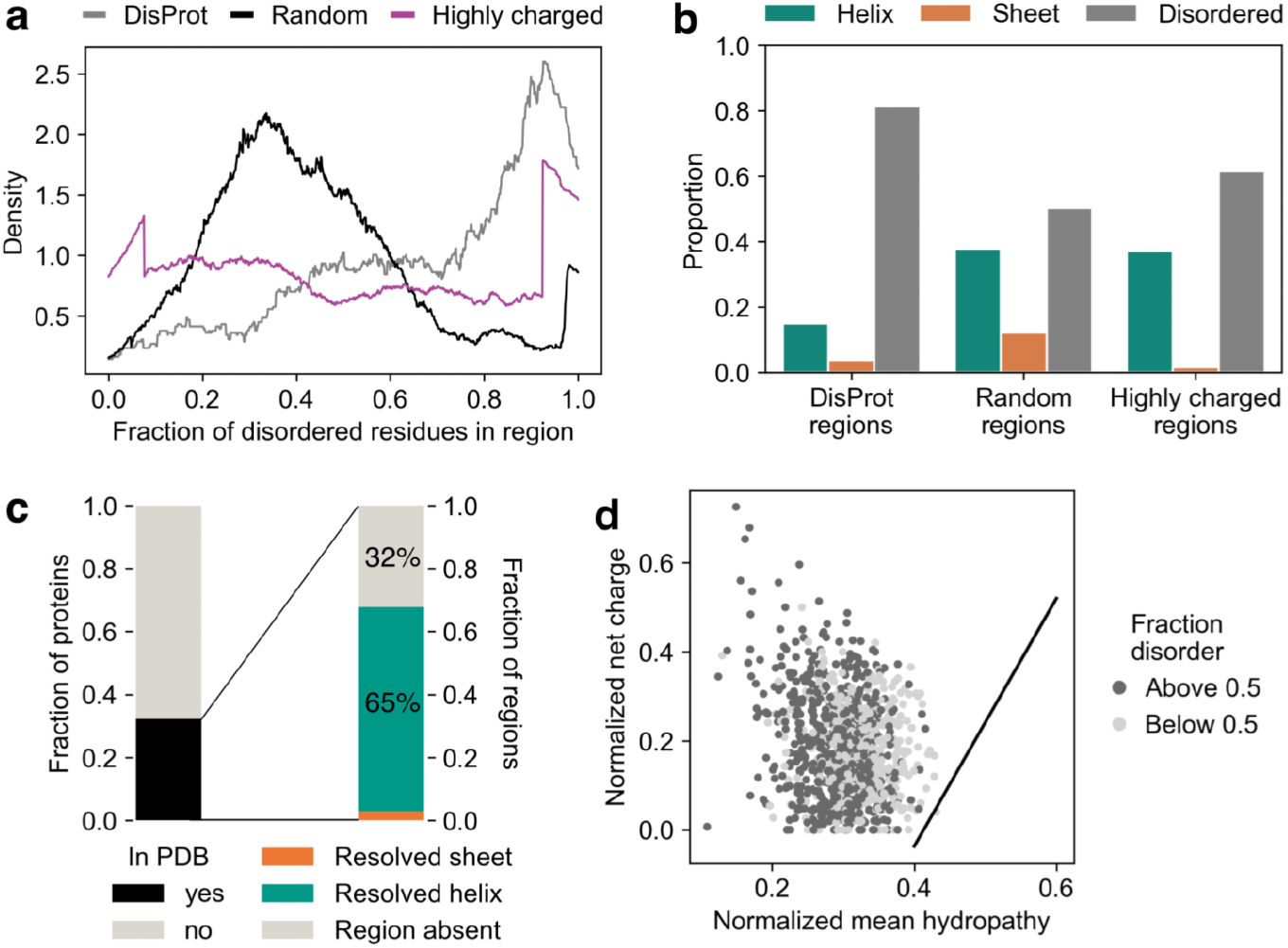
Secondry structure is pervaslve in highly charged, low complexity regions. **a** the distribution of the fraction of a region predicted to be disordered with AlphaFold, See Methods for secondary structure scoring. **b**. The number of predicted helical, sheet, and disordered (which includes coil; see Methods) residues for the highly charged regions, randomly-selected regions, and experimentally verified disordered regions from DisProt. **c** (Lef) The proportion of the top 200 most structured regions for which the protein that contains them is in the PDB. (Right) For those that are in the PDB, the proportion of regions that is a helix, sheet, or missing. **d**. An Uversky plot of the highly charged regions. The dividing line is from [2].

To validate the predictions, we examined the subset of proteins which have empirically determined structures. We searched the Protein Data Bank (PDB) for the 200 regions with the highest predicted fraction of structured residues (>92% predicted structured, 19% of the total regions). 27% of the proteins in this subset could be found in the PDB and, of those that were found, 42 (68%) had the highly charged region resolved. In all of these cases but two (3%), the region predicted to be a helix was experimentally determined to be a helix (Figure 2c). That is, where experimental validation is available for these regions, odds are better than 20:1 that the helical prediction will be confirmed by experimental data.

In roughly a third of cases the region is absent from the PDB, and because disordered structures frequently evade structural resolution, it is possible that these regions are disordered under some conditions. In particular, solvent conditions (e.g., pH) and sequence context could modulate the net charge on each amino acid, altering the propensity for structure. However, given that these local effects are challenging to predict [38], and in principle could both promote or inhibit structure formation, we consider our estimate a reasonable lower bound on the propensity for helix formation in these regions. It is also possible that the helical region is stable within a disordered element, like a pipe on a jump rope, or that only a portion of the protein lacking the highly charged region was expressed and characterized. From our analysis, we conclude that there is no evidence to suggest structural predictions are inaccurate for these regions, and we confirm the presence of many highly charged helices in experimental data.

The presence of substantial helical structure in these highly charged, low-hydrophobicity regions raised questions as to how these regions would be scored by the metrics which initially established connections between disorder and these sequence features. As a set, these regions contradict the argument that large amounts of uncompensated charge predicts disorder, captured in the popular charge/hydropathy or Uversky plot [2]. We created a Uversky plot of the highly charged regions, and all but three fell above the dividing line into the “natively unfolded” region (Figure 2d). Thus classic methods for determining whether a region is ordered or disordered have virtually no predictive power for regions of this composition—a surprising result given that these regions would appear perfectly suited for such a heuristic.

More recently developed metrics have been used to assign biophysical properties to highly charged regions. In particular, connections have been made between the patterning of charges and the predicted radius of gyration (R_g_, a measure of compaction) [3,9]. R_g_ is used to characterize ensembles of disordered conformations; the implicit argument appears to be that because most IDRs are polyampholytes, analysis of polyampholyte conformations can be carried out productively without considering structured conformations. Yet in specific cases, the very polyampholyte sequences being assigned to various disordered conformational states by computational analysis due to their charge patterning [3] are known to be helical—such as the (EEEKKK)n and (EEEEKKKK)n polymers [31,38,39]. Remarkably, this is more than a mere conceptual curiosity: our set of highly charged sequences in budding yeast contains the sequence KKKKEEEEKKKKEEEEKKKKEEEEKKKKEEEEKKKKEEEEKKKKEEEEKKKQEEEEKKKKEEE EKKKQ in the protein Mnn4, a region which is, with modifications, conserved in other fungal species (Fig S2c). Such sequences and their relatively well-studied biophysical behavior offer additional evidence of the importance of considering helical structure in biologically relevant highly charged regions.

These particular sequences show the hallmarks of so-called single alpha helices (SAH), helices of length 25–200 residues frequently, though not exclusively, formed by (E_4_K_4_)_n_ repeats [9,33,34]. Below, we establish the evolutionary conservation of these regions, suggesting their structure, as well as high charge density, are likely to confer a fitness benefit.

### An ancient translation initiation factor contains a conserved highly charged helix with sequence properties similar to an IDR

To more deeply investigate the sequence and structural properties of highly charged regions, we focused on a specific example where a region predicted to be a helix by AlphaFold had a solved empirical structure for comparison. We chose the broadly conserved eukaryotic translation initiation factor eIF3A (Rpg1 in *S. cerevisiae*) in which we identified several highly charged regions. One such region was predicted to be almost entirely helical, which is confirmed in the cryo-electron microscopy (cryoEM) structure (Figure 3b) [40].

**Fig. 3.**
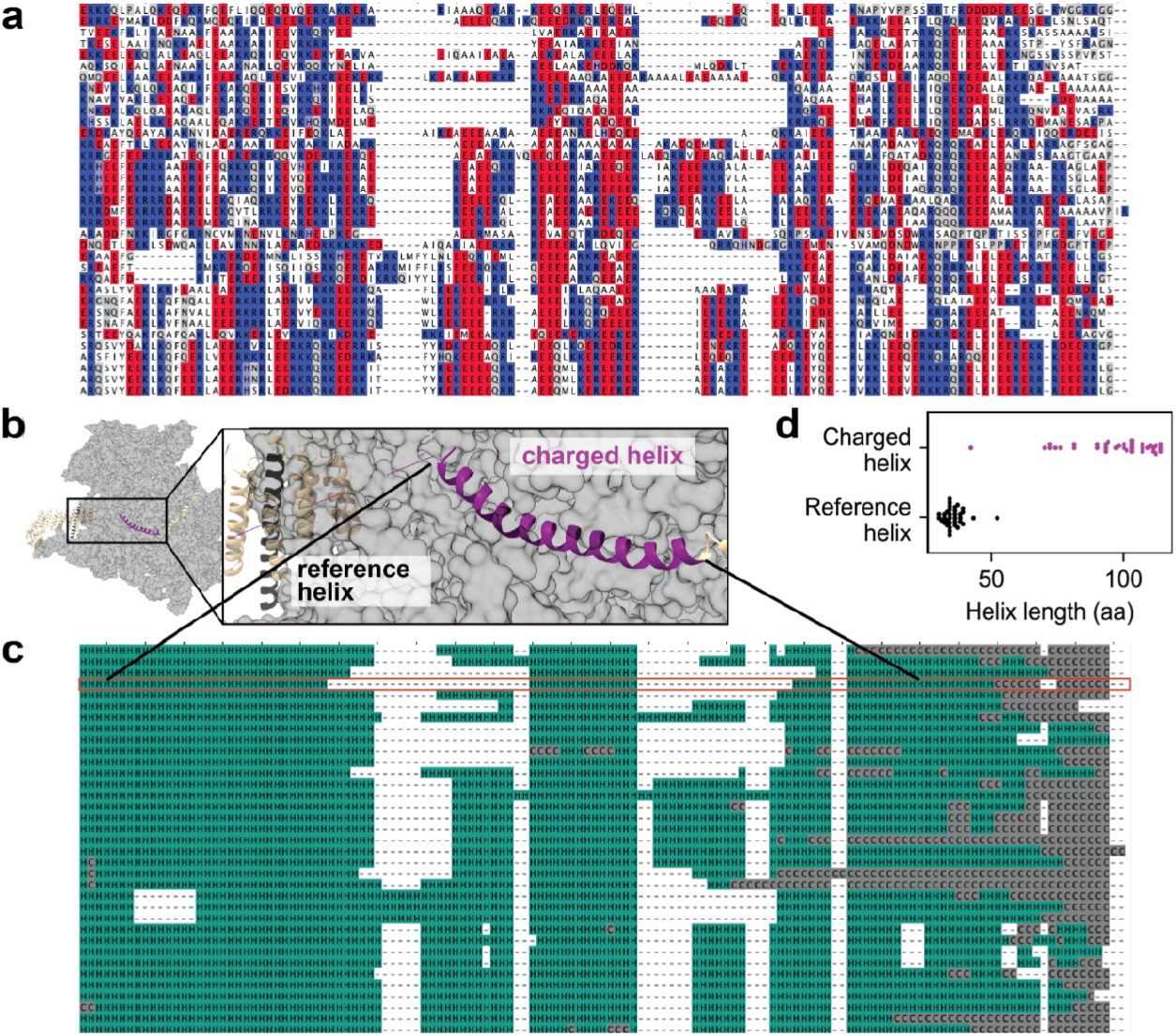
elF3A, an essential eukaryotic translation initiation factor, contains a conserved, highly charged helix that varies in length but not in secondary structure. **a** Alignment of the elF3A highly charged region (orthologs from all distantly-related species with predicted proteomes in AlphaFold) with negatively charged residues colored red, positively charged residues colored blue, and gaps and all other amino acids in white. Although the highly charged nature of the region is conserved, the sequence itself is variable. **b** Representative image of the cryoEM structure of yeast elF3A [40] with the highly charged region (resolved as a helix) shown in purple, and a reference helix shown in black. **c** Alignment of elF3A (same as in a) colored by the secondary structure predicted by AlphaFold; *S. cerevisiae* sequence is highlighted in red. Note that the highly charged region is predicted to be helical in every species represented. **d** Despite strong secondary structure conservation, the length of the highly charged helix varies significantly more than a reference helix from the same protein.

When we created a sequence alignment of all the homologous proteins for which a structure had been predicted from AlphaFold, we found significant variation in both the length and the sequence of the region (Figure 3a). Such variability is typical in disordered low-complexity regions and seen as the accumulation of many insertions and deletions in a multiple sequence alignment, but we were surprised to see this variation because the yeast version of this sequence was structured. To determine whether the homologous sequences were likely to be structured as well, we used DSSP (implemented in the MDTraj package for Python [41]) to classify the secondary structure predicted by AlphaFold and projected the predictions into alignment space (Figure 3c). Despite the lack of conservation at the sequence level, the helical nature of the region was conserved across all the homologs. However, the length of the helix varied significantly: the coefficient of variation of the highly charged helix was 0.17, compared to 0.11 for a reference helix located elsewhere in the same protein (Figure 3d).

This analysis demonstrates that a region with all the features typically associated with IDRs (high length variation as indicated by gaps in the alignment, poor sequence conservation, low complexity, low hydrophobicity) can be associated with a charged region that is in fact structured and retains this structure across evolutionary time.

### Helical highly charged regions can be predicted from amino acid composition

Given the prevalence of structure in the highly charged regions that we detected, and the failure of existing composition-based heuristics to discriminate between the two categories (disordered and helical), we set out to determine if there were alternative simple heuristics that are effective. We stress that our goal is to develop insight into the factors that determine disorder versus structure, not to replace sophisticated software [32,33,42,43]. Using the proteome-wide predictions of structure from AlphaFold, we created a dataset of regions which were predicted to be either completely disordered or completely helical (13,437 sequences from 63.6% of the proteome). On a Uversky plot, the helices and IDRs drawn from the yeast proteome, like the highly charged regions, could not be distinguished on the basis of normalized mean hydropathy and net charge (Figure 4a).

**Fig. 4.**
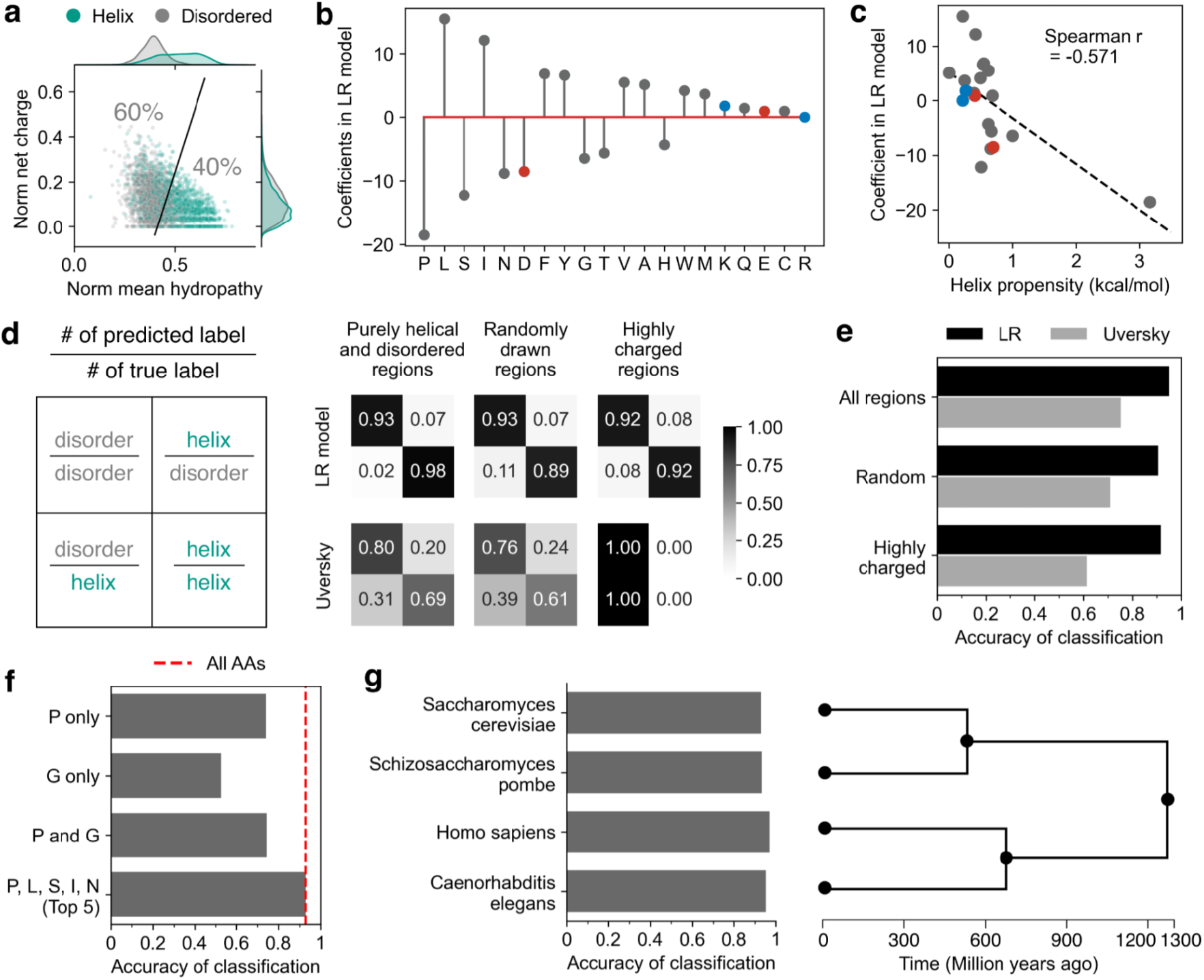
Helical regions can be predicted from composition. **a** Uversky plot of all regions used to train the LR model. The marginals of the distribution are shown on the plot border. **b** The coefficients of the logistic regression (LR) model which predicts whether a region is helical or disordered on the basis of amino acid composition. The model was trained on purely helical and disordered regions (predicted by AlphaFold) selected from the *S. cerevisiae* proteome. Amino acids with a positive coefficient are correlated with helices, those with a negative value are correlated with disordered regions. **c** The helix propensity from [44] plotted against the LR model coefficients. **d** Accuracy of both the LR model (top) and the Uversky dividing line (bottom, from [2]) on purely helical and disordered regions (held out data from the training set, n=3360; left), randomly-drawn regions, which are predicted by AlphaFold to be majority (but not completely) helical or disordered (n=3405; center), and the highly charged regions (n=681; right). **e** Summarized accuracy for all categories in **d. f** Summarized accuracy of the LR model with only a subset of coefficients, **g** (Left) Accuracy of the LR model prediction of regions from other organisms. (Right) Timetree showing the evolutionary divergence of the organisms.

Using this dataset, we built a logistic regression model which classified regions as helical or disordered on the basis of their amino acid composition (see Methods for model details). The coefficients of this logistic regression (LR) model are shown in Figure 4b; as expected, residues known to affect helical character such as proline and glycine have a large regression coefficient, indicating that their presence is highly predictive. More generally, the coefficients from the model are inversely correlated with the individual amino acid helix propensity [44] (Figure 4c).

We assessed the accuracy of the LR model in several ways. First, we calculated the rate of true and false positives and negatives (Figure 4d) for several classes of sequences. The LR model performed extremely well, correctly identifying both helical and disordered regions in the testing data (25% held out from the original dataset) with an accuracy of 92.5% (Figure 4e). We also assessed its performance on a new set of randomly selected regions which were predicted by AlphaFold to contain both helical and disordered character; the LR model predicted the dominant structural feature from composition alone 86.9% of the time (Figure 4e). Finally, we predicted the highly charged regions, where the LR model performed with an overall accuracy of 90.8% (Figure 4e). Most of this accuracy can be captured using only the top five coefficients of the model (Figure 4f).

We also used the LR model built from AlphaFold predictions to score a dataset of PDB structures with secondary structure annotation (see Methods); the model performed with an accuracy of ∼90% on these experimentally determined structures (Figure S4c). To see whether there were systematic differences in the relationship between amino acid composition and secondary structure when using real versus predicted structures, we also created a second LR model trained on purely helical or disordered sequences from this PDB-derived dataset and compared it to the original LR model (Figure S4a, S4b). The coefficients of the two models are highly correlated, with the interesting exceptions of the two helix-breakers P (which has the same order of importance in both models but a much larger magnitude coefficient in the PDB LR model) and G (which has a much higher relative importance in the PDB LR model).

To put our results in context and understand the breakdown of existing heuristics, we compared the accuracy of the LR model to the accuracy of the charge/hydropathy or Uversky model. As expected, the Uversky model performed better than chance but worse than the LR model on the same sets of randomly drawn regions, but was completely non-predictive for highly charged regions (Figure 4e). This reinforces the idea that normalized net charge is not predictive of disorder (see marginal distributions in the right hand side of Figure 4c), at least for regions with this length distribution. In sum, virtually all the predictive power in the Uversky charge/hydropathy heuristic comes from hydropathy.

Finally, we were curious whether our model was specific to amino acid usage in yeast, or if it could be extended to other proteomes. Using three other proteomes for which structures have been predicted, *Schizosaccharomyces pombe, Caenorhabditis elegans*, and *Homo sapiens*, we performed the same procedure of random region selection, labeling using the AlphaFold predictions and confidences, and classification with the LR model trained on AlphaFold predictions of yeast regions. The prediction accuracy was nearly identical to or slightly higher than the *S. cerevisiae* accuracy (Figure 4g). This simple model based only on the composition of a region is sufficient to predict helical or disordered character in proteomes that diverged over a billion years ago.

### Highly charged regions are evolutionarily conserved

The unique evolutionary signatures suggested by our analysis of eIF3A, coupled with the consistency in predictive power of the LR model across vast evolutionary distances, led us to broader questions about the conservation of the regions we had identified. To what extent do highly charged regions retain their sequence properties and structure as organisms evolve?

To address this question, we turned to AYbRAH, a curated database of protein homologs and paralogs in 33 fungal species spanning 600 million years of evolution [45]. This dataset combines automated homology detection and manual curation to achieve high-confidence predictions of highly diverged orthologs.

First, we quantified the sequence conservation of the regions in question by examining their alignments and calculating both the frequency of gaps and the sequence divergence (average position-wise entropy). Sequences with high insertion and deletion rates, a known feature of IDRs, will have higher alignment gap frequencies. Those with a high point mutation rate will have high divergence. A low value of both metrics indicates sequence conservation. We calculated these values for all the highly charged regions, and compared them to length-matched, randomly drawn regions from the rest of the same proteins. We found that the charged regions have significantly more gaps (*P*<0.001, Mann-Whitney U test) and are more divergent (*P*<0.001, Mann-Whitney U test) than the proteins in which they are found (Figure 5a). We compared these distributions to the same values calculated for experimentally verified IDRs from DisProt, and found that although the charged regions have similar gap frequency to IDRs, they are even more divergent at the sequence level. Thus simply viewing an alignment of these LCRs, without using a secondary structure prediction algorithm, one might conclude that they are disordered.

**Fig. 5.**
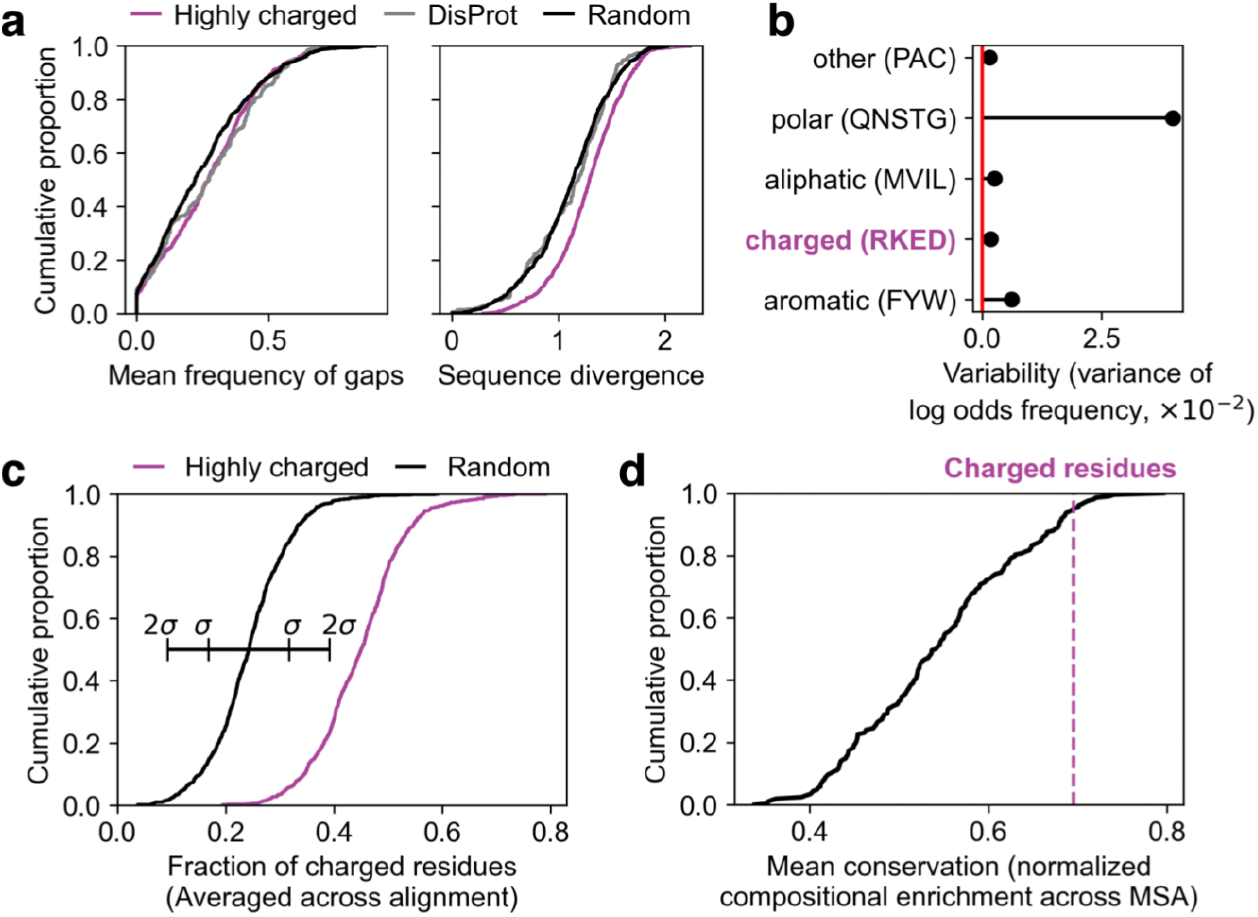
Highly charged regions are evolutionarily conserved. **a** The distribution of gap frequencies (left) and average position-wise entropy (right) summarizing multiple sequence alignments (MSAs) of the highly charged regions (purple), randomly drawn regions from the rest of the proteins that contain them (black), and IDRs from DisProt (gray). **b** Summary of the variance (in log odds space) of usage of categories of amino acids for all proteomes in the AYbRAH database. **c** FCR values, averaged across AYbRAH alignments, for the highly charged regions identified in the *S. cerevisiae* proteome and length-matched randomly-drawn regions and their associated AYbRAH MSA. **d** The average compositional conservation of regions enriched for all sets of four amino acids with the same total frequency as the charged amino acids, plotted as a CDF. Higher values indicate less drift, and lower values indicate more drift (regression to the proteome average).

Despite this apparent lack of conservation, we were curious whether any aspects beyond sequence of the highly charged regions were conserved. These regions were identified because of their unique sequence composition, so we tested the degree to which they retained this composition over evolutionary time. To first determine the expected compositional variation, we measured the variation in the total proportion of each amino acid across the species represented in the AYbRAH database (Figure S5a). We found that as a group, the charged amino acids had very little variation in proportion of usage (Figure 5b). Consistent with selection, a high fraction of charged residues was preserved across species and substantially differed from randomly drawn regions in the same species (Figure 5c).

The distribution in Figure 5c contains some regions that on average fall below the threshold of 0.4 FCR that we established for the original search in the yeast proteome; this is not surprising given that all sequences are subject to drift, which pulls them towards the proteome average for any given trait unless selection intervenes. Therefore, we created a method to quantify drift in charged regions relative to other compositionally extreme regions.

We first identified regions in the yeast proteome enriched for all groups of four amino acids with a combined frequency within +/–0.01 of the combined frequency of the charged amino acids (0.233). For each of the 209 datasets, we calculated the mean proportion of the amino acids in question for each identified region across the AYbRAH alignment (note that FCR is a special case of this property where the four amino acids in question are glutamate, aspartate, lysine, and arginine) If the composition (enrichment of the four amino acids in question) is conserved, we should expect that the mean enrichment score across regions and alignments should be close to the mean of the original enriched dataset (close to or higher than the 0.4 threshold). In contrast, if the property is not conserved, we should expect this enrichment score to be close to the proteome average. To compare directly between datasets, we scaled this enrichment score to a unit scale between an effective 0 (the proteome average), and 1 (the median of the enriched regions detected from the *S. cerevisiae* proteome). Sets of four with a conservation score close to 1 are highly conserved for that property, while those that are close to 0 have experienced high levels of drift, indicating that they are not conserved. We find that the set of four charged amino acids falls within the top 5% of these scores (Figure 5d). From this analysis we conclude that in the highly charged regions, the charge density is extremely well-conserved.

## Discussion

To understand the biology of proteins and their subdomains, heuristics are almost inevitably used: comparison to other proteins to infer similarity by homology, motif identification to predict binding partners, and so on. In the case of highly charged protein regions, several heuristics appear to converge on the conclusion that such regions are overwhelmingly likely to be disordered. By virtue of their strongly biased sequence composition, they tend to fall into the class of low-complexity sequences associated with lack of stable structure [46]; they tolerate insertion/deletion events at higher rates than typical well-folded sequences; their high charge favors interactions with solvent, and low hydrophobicity suggests the absence of a solvent-protected hydrophobic core. Consistent with this, many analyses of such regions proceed as though structure can be mostly or completely ignored.

Here, we have shown that naturally occurring highly charged regions are predicted to adopt helical structure to a degree which cannot be neglected — ∼40% in a proteome-wide analysis — and that these predictions are validated by existing experimental data for both structured and disordered sequences. Moreover, we show that all these heuristic signals of disorder are in fact compatible with fully structured polypeptides: extended charged helices which have no hydrophobic core, grow and shrink in length over evolutionary time, interact with solvent on all sides, and form from sequences of two or even one type of amino acid (Fig. 6a). Together, our results indicate that understanding the biology of highly charged sequences requires integrating insights from both structural and disorder-based approaches.

**Fig. 6.**
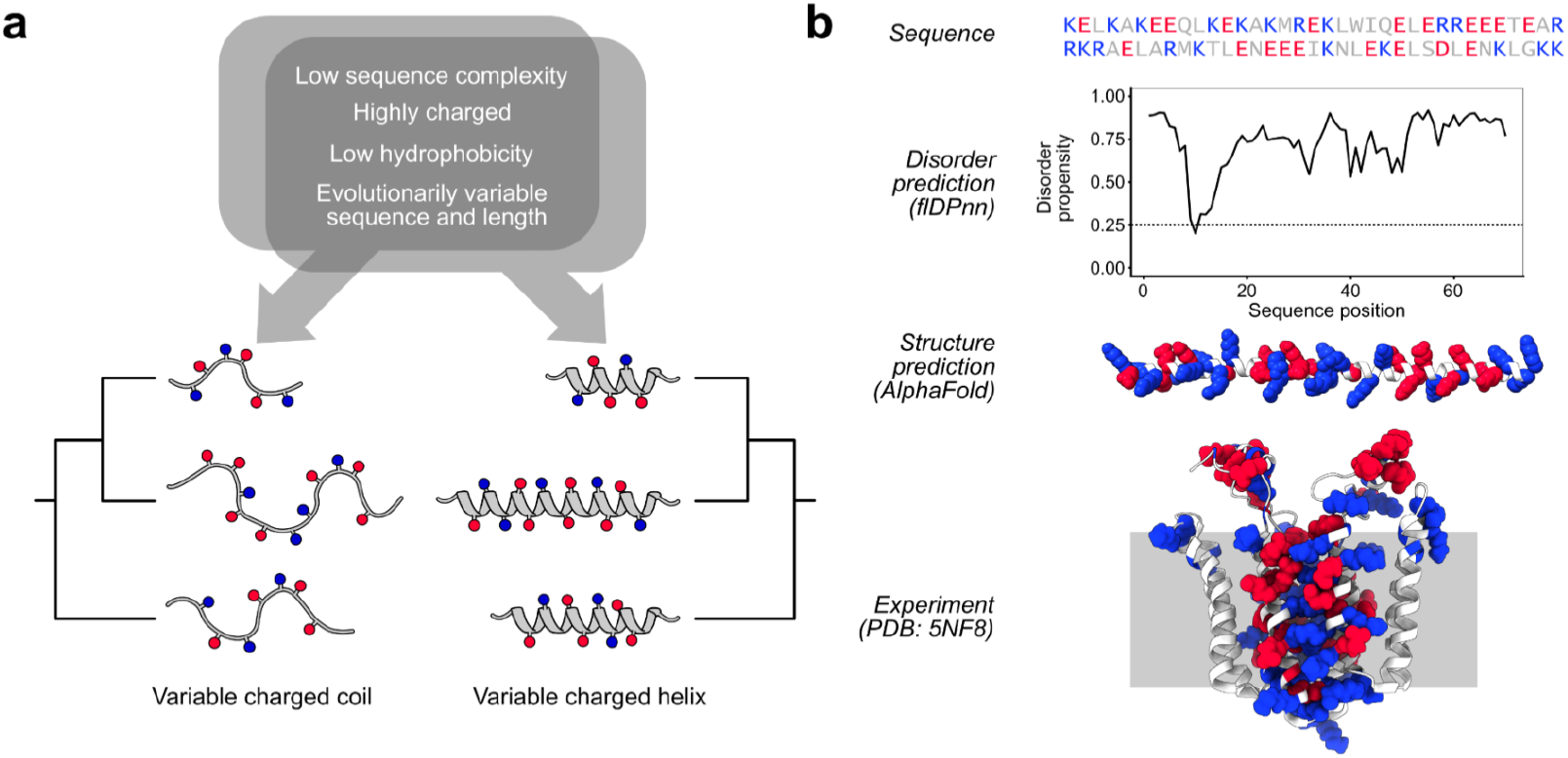
Rethinking assumptions of disorder in highly charged regions. **a** The same set of predictors that have been associated with disordered regions turn out to also be compatible with a fully structured helical region. **b** Present leading-edge methods for disorder and structure prediction disagree completely and also fail to capture experimental reality. The highly charged region of Rcf1 is alternatively predicted to be near-completely disordered or completely helical; experimentally and biologically, it forms most of the membrane-spanning helices and exposed loops in this dimeric mitochondrial inner membrane protein.

A consequence of these results is that they upend multiple well-established heuristics for determining how to think about, and study, a sequence’s biological behaviors. The shortcut that charge and hydrophobicity can serve as accurate dimensions for separating structured from disordered sequences, powerfully demonstrated using limited data available at the time [2], does not work for highly charged, low-hydrophobicity sequences. Although many sophisticated methods for detecting disorder or single helices have been developed [32,33,42,43,47], the assumption that, because many disordered sequences are polyampholytes, other polyampholyte sequences can reasonably be treated as if disordered persists in modern work [3]. We emphasize that our results say nothing about the utility of further results built on the assumption of disorder, conditioned on its accuracy. And we further stress that considerable work may properly focus only on disordered sequences with no claims regarding the assessment of disorder. Nevertheless, as for the example of (E_4_K_4_)_n_ polymers, it is straightforward to find examples in which sequences known to have well-defined structure are treated as if they did not, evidence for the undue influence of improper heuristics.

Given these results, it might seem inevitable that sophisticated structure-prediction methods would be required to more accurately discern whether particular highly charged sequences adopt a helical conformation. However, we introduce a simple amino acid composition-based classifier—logistic regression with as few as five inputs—which can predict structure (or its absence) with accuracy above 90%. This model is trained on biologically occurring sequences, a tiny and profoundly biased subset of protein sequence space, such that we do not expect its performance to carry over to arbitrary sequences. Still, as a heuristic method implemented with a handful of numbers, it balances simplicity and accuracy (particularly over the charge/hydropathy heuristic) in a way which is practically useful in diagnosing structure for charged sequences.

The notion that intrinsically disordered proteins or regions sometimes adopt structure is well-understood [34], particularly in the case of folding upon binding [48]. Because of this distinction between conformations in isolation versus when bound to a partner, structures in the PDB may tell only a portion of the story. Similarly, AlphaFold specifically predicts structures most likely to appear in the PDB [32], rather than, for example, conformations which are occupied most of the time in the biological context. To the extent that our results depend on these resources, they similarly remain inapplicable to questions about the broader conformational ensembles that highly charged sequences may sample. But how stable or frequently adopted might these structures be?

From the perspective of evolutionary conservation, even a conformation which is occupied for a tiny fraction of the time may impose dominant constraints on a sequence, if this conformation contributes to organism fitness. To the extent that we wish to understand the relationship between sequence and biological function, this potential for rare conformations to dictate function may permit most conformations in the ensemble to be neglected—much as recognition of folding upon binding for a disordered region may properly focus attention on the bound state, even if it is fleeting. In the case of highly charged regions which must adopt helical conformations to carry out their functions, certain near-absolute constraints must be satisfied; no matter how unstable the helix, a proline kink in the backbone cannot be straightened, and so depletion of proline from these regions provides an additional signal. On the other hand, presence of a proline powerfully indicates that a straight helix cannot form and is therefore unlikely to be the functional conformation, no matter what other sequence signals exist and no matter how fleeting the helix state is proposed to be.

Even the best methods for predicting structure and for predicting disorder can disagree and fail to capture experimental reality. Consider the highly charged region of the yeast protein Rcf1 (Fig. 6b). A top-ranked modern disorder predictor, flDPnn [42,47], predicts this region to be entirely disordered. AlphaFold predicts it to be entirely helical with high confidence. Neither captures reality: experimentally, this region forms most of a dimeric five-pass transmembrane protein in which charged residues, exposed on stable helices, form dimer-stabilizing salt bridges through the mitochondrial inner membrane [49] (Fig. 6b). To the extent that cases like this closing example persist, the challenges we identify here remain open.

Broadly, while our results uncover previously overlooked structure in highly charged regions, the dual challenges of determining the biologically active configurations of these sequences, and of determining the statistical features of the conformational ensembles they occupy, remain open. Rather than looking at such sequences through the lens of disorder, it appears that both lenses—structure and disorder—will be needed to give the proper depth of focus.

## Methods

### Extraction of highly charged regions from the yeast proteome

The S288C reference genome was obtained from the Saccharomyces Genome Database (SGD). For each gene in the reference genome, we first computed fractional charge as a moving average across its sequence using a window size of 12 residues and a triangular weight, where the highest weight was assigned to the middle region of each window. We then searched for highly charged regions in each sequence based on a fractional charge threshold of 0.4 and tolerance of 10 residues. Tolerance refers to the maximum number of residues that we allow to have moving average values below the fractional charge threshold before terminating the region. This tolerance allows for transient deviations from high charge and prevents fragmenting highly charged regions with small insertions of uncharged amino acids. We extracted regions that were longer than a given minimum region length of 30 residues, then trimmed any remaining uncharged residues from the N and C terminal ends of the sequences (these result from the triangular weighting scheme and the tolerance).

### Calculating sequence complexity

Sequence complexity was calculated according to Ref. 9 using the following equation:

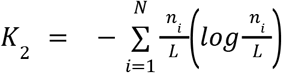

where *K*_2_ is the unnormalized complexity, *N* is the number of possible residues (in this case the 20 natural amino acids), and *n*_*i*_ is the number of each residue in the sequence, which has length

*L*. This value is normalized to the “entropy of the language” (e.g., the yeast proteome), such that a sequence with compositional properties exactly equal to the average frequencies will have a complexity of 1.

The entropy of the language is calculated using the equation

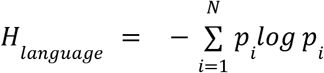

where *p*_*i*_ is the frequency of letter *i* in the reference. We used two different languages as references; the first is the yeast proteome (so each *p*_*i*_ represents the average frequency of that amino acid in the proteome). We also used a modified reference enriched for charged residues; each of the amino acids lysine, arginine, aspartate, and glutamate had a frequency of 0.125; the remaining frequency (0.5) was distributed evenly among the other amino acids.

### Generation of null distribution for amino acid usage

We counted the number of occurrences of adenine (A), thymine (T), cytosine (C), and guanine (G) in the DNA sequences of all open reading frames in *S. cerevisiae*. The expected frequency of each codon was computed as the product of the frequencies of all nucleotides that appear in that codon and the expected frequency of each amino acid was computed as the sum of the expected frequencies of all codons specifying that amino acid.

### Extraction of AlphaFold data

We used proteome-wide structure predictions from AlphaFold to analyze the structure of the regions we identified with high proportions of charged residues. We downloaded the structures for all *S. cerevisiae* proteins from the AlphaFold website (https://alphafold.ebi.ac.uk/download#proteomes-section) [32]. We read the PDB files into Python and used DSSP implemented in MDTraj [41] to score secondary structure. We used custom Python functions to extract the confidence scores from the predicted structure file for each protein.

### Construction of a logistic regression (LR) model to predict secondary structure

To classify the secondary structure of a region on the basis of composition, we first constructed a training dataset built from AlphaFold structures. We classified all residues in all structures as either helical (classified as helical by DSSP and with a pLDDT score above 70), disordered (classified as coil by DSSP and with a pLDDT score above 70 or any residue with a pLDDT score less than 50) [37], sheet (classified as sheet by DSSP and with a pLDDT score above 70), or other. The pLDDT cutoff of 70 marks was chosen because this value was used by the creators of AlphaFold to distinguish between “Confident” and “Low” model confidence.

To generate training and testing data for the LR model, we exhaustively searched for purely helical and disordered regions by identifying regions that were greater than 25 amino acids long and only contained either helical or disordered residues. We extracted 6882 helical regions and 6366 disordered regions from the *S. cerevisiae* proteome. We randomly selected 75% of these regions as training data and 25% of the regions as testing data and built a LR model using amino acid composition as predictors. The LR model is built using the scikit-learn package in Python.

We also constructed a LR model based on empirical structures from the PDB; secondary structure and disorder were annotated on a per-residue basis from the experimental 3D structure by the PDB and were obtained by request [50] (see their Methods for details). The final dataset contained 64,804 regions greater than 25 amino acids long and consisting solely of either helix or disorder; the regions were approximately equally split between helical and disordered. Otherwise, the procedure was identical to that used for the AlphaFold predictions.

### Classifying regions and computing model accuracy

We used the per-residue secondary structure classifications described above to score the highly charged regions: regions with more than 60% helical or disordered residues were labeled as their dominant type, and all others were labeled as “intermediate.” This is the “true” label. Out of all the regions, 34% were labeled as disordered and 31% were labeled as helical. We then used the logistic regression model to classify these regions on the basis of their amino acid composition. The true labels were used to compute the false negative and positive rates, and the overall model accuracy.

To directly compare to this dataset, we also randomly drew regions from the AlphaFold dataset with the same scoring conditions as we used for the highly charged regions (>60% helical or disordered), scored the region using the model, and computed accuracy in the same way.

### Evolutionary analysis: sequence properties

We used custom Python scripts to extract multiple sequence alignments (MSAs) for all proteins in the AYbRAH alignment [45]. A region was considered “present” in a protein in the MSA if the region which aligned to the *S. cerevisiae* sequence that we identified as highly charged contained at least 30 amino acids (the same length minimum length as was required for a region detected by the algorithm).

To compute alignment quality, the longest and shortest sequences in each alignment were removed, and any resulting columns containing only gaps (represented by the symbol “-” in the alignment) were removed. We then quantified the frequency of gaps as well as sequence column-wise entropy, which we refer to as “sequence divergence.” The mean frequency of gaps was computed as the number of “-” characters divided by the total number of characters across all sequences in an alignment. Sequence divergence was computed by summarizing the frequency of amino acids in each column as a one-dimensional probability distribution and calculating the Shannon entropy of that distribution.

### Evolutionary analysis: compositional drift

To test compositional drift for regions enriched in selected amino acids, we identified all unique sets of four amino acids (excluding the charged amino acids) with the same combined frequency as the four charged amino acids. We modified the charged region algorithm to detect regions enriched for these sets of four. We used an enrichment threshold of 0.35 (35% of the region composed of the amino acids in question), minimum length of 30 amino acids, and a tolerance of 15; these parameters were selected to yield datasets that most closely matched the number of hits and length distribution of the original dataset (enrichment for E, D, R, and K). For each of these datasets, we calculated the conservation by averaging the enrichment score (percent composition of the specific amino acids) across the alignment in each region, taking the mean of that distribution, and scaling it between the *S. cerevisiae* proteome average (0) and the average enrichment of the hits detected by the algorithm (1). This allowed us to compare datasets directly: sets of amino acids with values close to 0 experienced enough drift that they approach the proteome average; those with values close to 1 stay far from the average and close to the (rescaled) enrichment threshold and thus are likely conserved.

### Analysis of the eIF3A charged helix

We used MUSCLE version 3.8.31 [51] with default parameters to generate an alignment of all the eIF3A homologs for which a structure was available on AlphaFold as of April 2022 (35 species of model organisms including bacteria, yeast, mold, rice, soybean, mouse, and human). This dataset was used in Figure 3.

A cryoEM structure of eIF3A in complex with the ribosome was used for structural analysis. The structure has PDB code 6ZCE (http://doi.org/10.2210/pdb6ZCE/pdb) [40].

Secondary structure was scored with DSSP as described above. The length of the charged helix was calculated by identifying the start of the highly charged region and then counting the amino acids until a run of more than three non-helix characters was encountered. A reference helix from earlier in the sequence was chosen as a comparison.

### Analysis of Rcf1

The 70-residue Rcf1 charged region (Fig. 6b) was used for disorder predictions with flDPnn [42] at http://biomine.cs.vcu.edu/servers/flDPnn/ with default settings, and for structure prediction with AlphaFold through ColabFold [43].

### Statistical Tests

All p-values were calculated with the Mann-Whitney U Test (Wilcoxon Rank Sum Test), either two-sided if no hypotheses were formed about the relationship between the two distributions or one-sided otherwise. The evaluation of these tests was done in Python and can be found (along with the data) in the Jupyter notebook for the relevant figure.

## Data Availability

Data used in this study are from publicly available datasets: AlphaFold protein structure prediction available at https://alphafold.ebi.ac.uk/download#proteomes-section, yeast proteome available from Saccharomyces Genome Database http://sgd-archive.yeastgenome.org/sequence/S288C_reference/orf_protein/, AYbRAH fungal ortholog database available at https://github.com/LMSE/aybrah, and DisProt yeast disordered regions https://www.disprot.org/browse?sort_field=disprot_id&sort_value=asc&page_size=20&page=0&release=current&show_ambiguous=true&show_obsolete=false&ncbi_taxon_id=559292. All additional data generated in this study are available at https://github.com/drummondlab/highly-charged-regions-2022.

## Code availability

All analyses and code used to generate the figures in this work can be found at https://github.com/drummondlab/highly-charged-regions-2022.

## Author contributions

D.A.D., A.R.D. and C.G.T. developed ideas and direction, R.W.P. and C.G.T. performed analyses, R.W.P., C.G.T. and D.A.D. made figures, and all authors contributed to the text.

Acknowledgments

C.G.T. is a Damon Runyon Postdoctoral Fellow supported by the Damon Runyon Cancer Research Foundation (DRG-2465-22). R.W.P. acknowledges support from the UChicago Biological Sciences Collegiate Division Summer Fellowship, Liew Family College Research Fellows Fund, and the UChicago Quantitative Biology Summer Fellowship. D.A.D. acknowledges support from the NIH (award numbers GM144278 and GM127406) and the US Army Research Office (W911NF-14-1-0411). A.R.D. acknowledges support from the NIH (award number R35 GM136381). The content is solely the responsibility of the authors and does not necessarily represent the official views of the NIH.

The authors thank Alex Holehouse for helpful discussions, and Alexander Cope for providing the structure-annotated PDB data.

## Additional information

Supplementary materials can be found in the accompanying PDF.

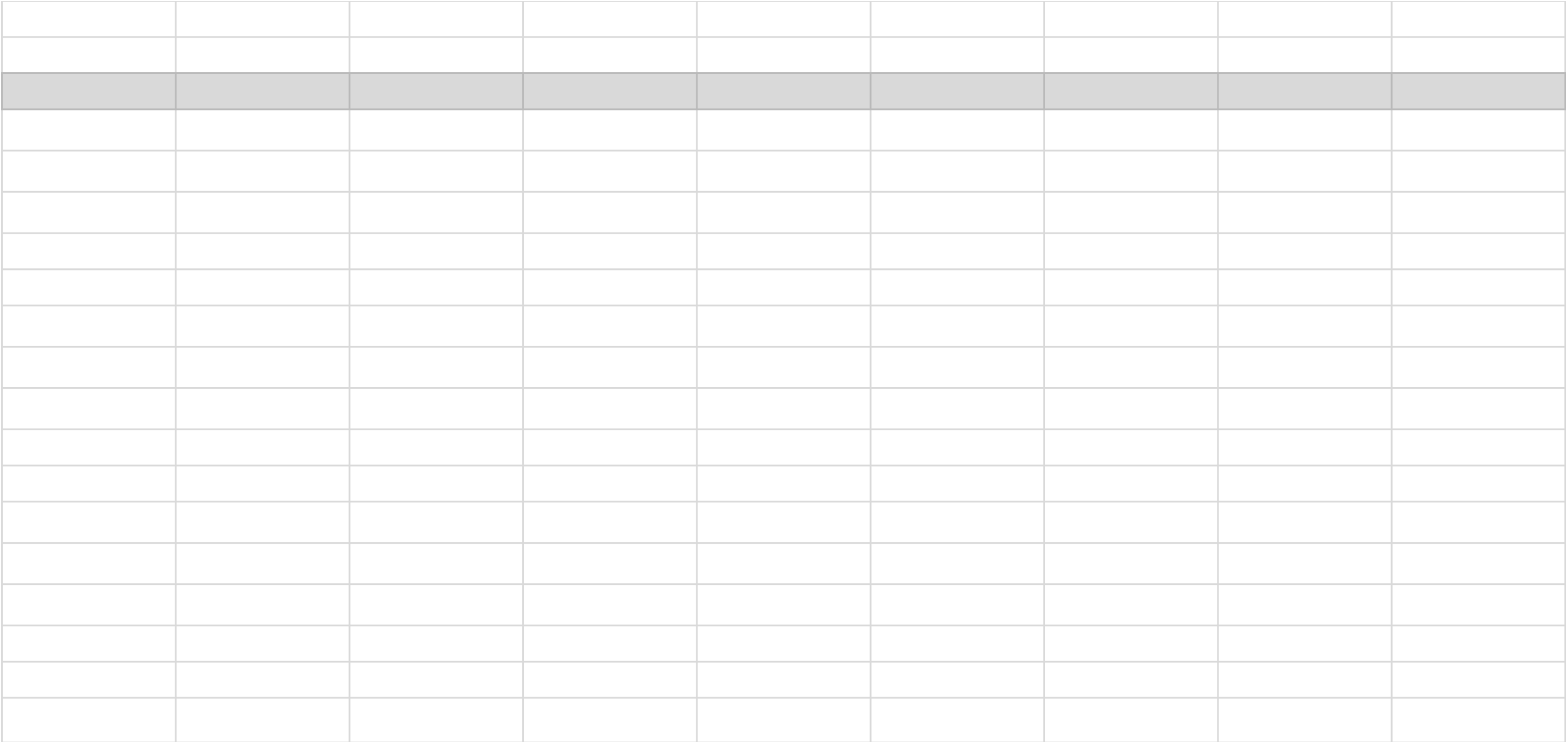

