## Supplemental Figures for "Pervasive, conserved secondary structure in highly charged protein regions"

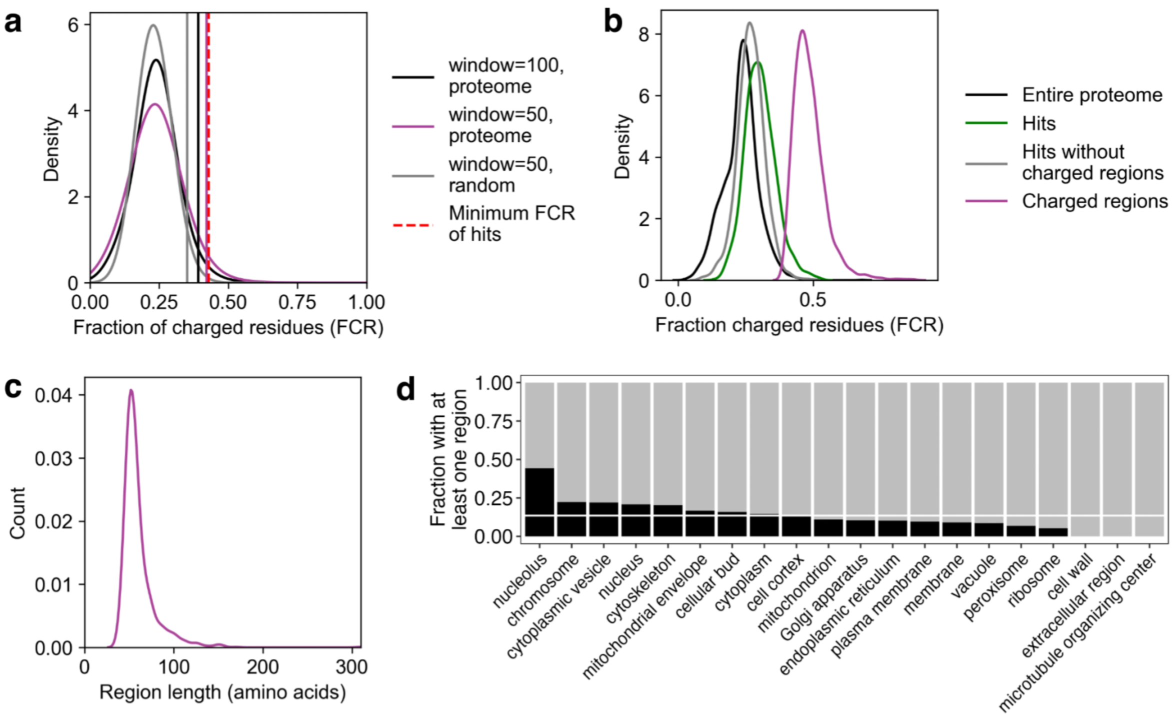


**Supplement to Fig.1: Regions high in charge are prevalent in the yeast proteome and are low complexity**
**a** The distribution of FCR values for randomly-drawn regions of different length in the yeast proteome (black, purple), and randomly generated regions using the average S. cerevisiae amino acid frequencies (gray). Vertical lines show two standard deviations above the mean. The minimum FCR of the highly charged regions after trimming is shown with a red dashed line. **b** The distribution of FCR values for the regions detected (purple), the proteins that contain them (green and gray), and the per-protein FCR for all yeast proteins (black). **c** Distribution of region lengths for highly charged regions. **d** Enrichment for proteins containing highly charged regions in different cell compartments. The proportion of the proteins in the category with at least one highly charged region is shown in black. The expected proportion (based on the number of unique genes detected by the algorithm and the total number of yeast proteins) is shown as a horizontal white line.

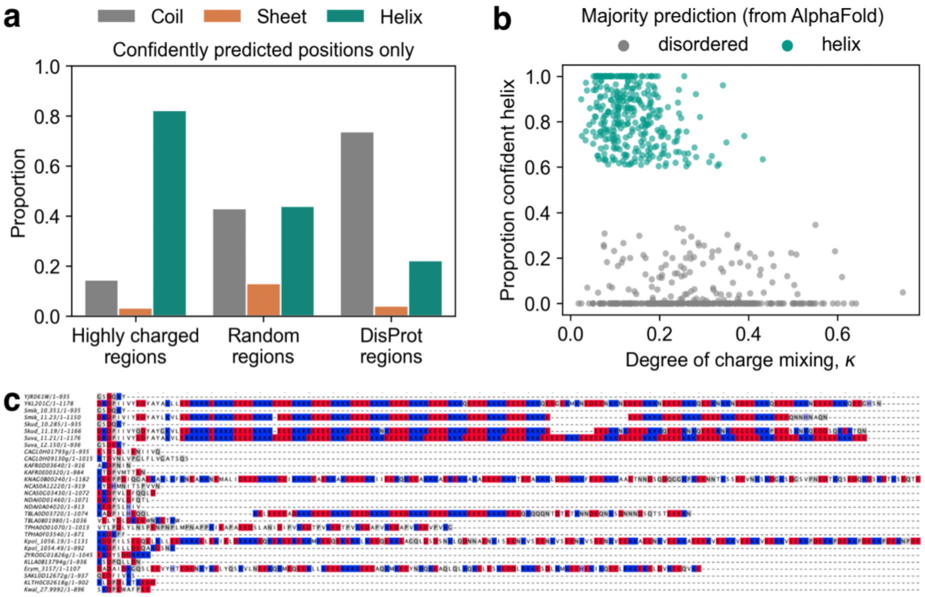


**Supplement to Fig.2: Secondary structure is pervasive in highly charged, low complexity regions**
**a** The frequency of (per-residue) secondary structure for all the confidently predicted (pLDDT > 70) residues from the AlphaFold structure for highly charged regions, length-matched random regions, and experimentally-verified disordered regions from DisProt. The majority of confidently prediction regions in disordered (DisProt) regions are coil. **b** The κ value of the region versus the proportion predicted helix for all regions that are more than 60% helix or disordered. **c** Alignment of MNN4 C-terminal region in fungal species in the AYbRAH database.

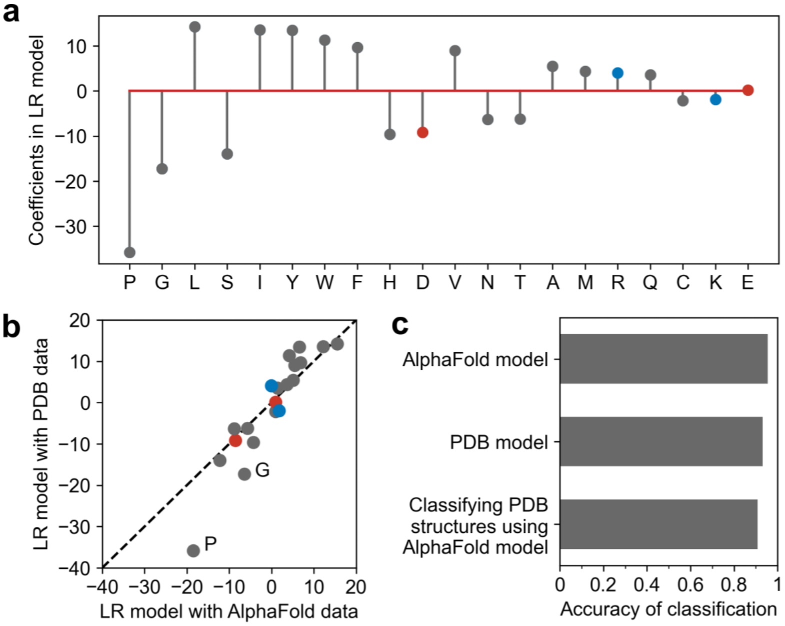


**Supplement to Fig.4 Helical regions can be predicted from composition**
**a** Coefficients from the LR model trained on helical and coil/disordered regions randomly drawn from the PDB. **b** Comparison between model coefficients in both the PDB and AlphaFold trained models. The importance of P and G varies the most between the two; P in magnitude and G in order of importance. **c** Accuracy of the AlphaFold and PDB trained models and the accuracy of the AlphaFold model when used to classify PDB structures.

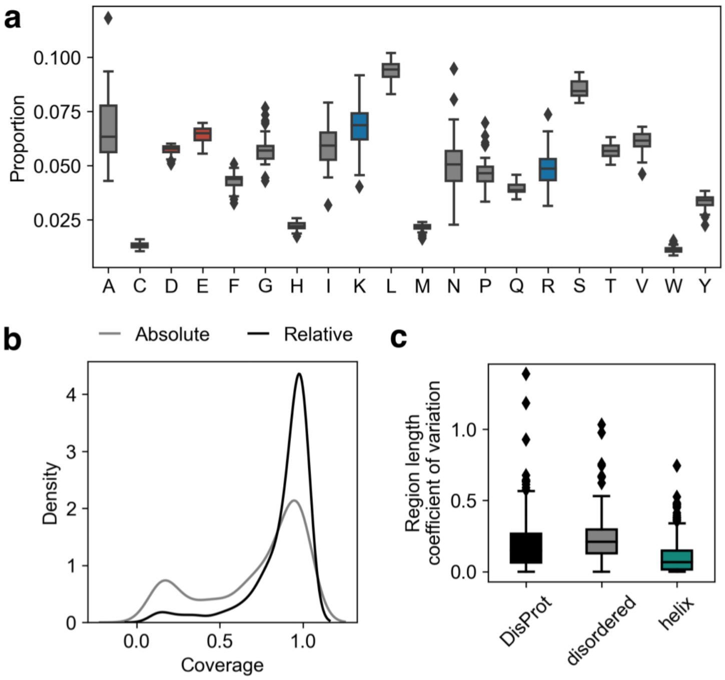


**Supplement to Fig.5: Highly charged regions are evolutionarily conserved**

**a** Variation in frequency for each individual amino acid. Distribution is the frequency of usage in each proteome in the AYbRAH database. **b** Absolute coverage (number of species in the AYbRAH alignment which contain non-gap characters in the detected highly charged region relative to the maximum number of species possible) and relative coverage (number of species in the AYbRAH alignment which have non-gap characters in the highly-charged region relative to the number of species with a homolog of the protein) for each protein with a detected highly charged region. **c** Length variation (between AYbRAH homologs) in regions labeled as helix (teal) or disordered (gray) based on their predicted structure from Alphafold and experimentally-verified yeast disordered regions from Disprot (black).
